# Complexity Measures of Psychotic Brain Activity in the Fmri Signal

**DOI:** 10.1101/2023.11.10.566647

**Authors:** Qiang Li, Masoud Seraji, Vince D Calhoun, Armin Iraji

## Abstract

When viewing the brain as a sophisticated, nonlinear dynamic system, employing complexity measures offers a valuable way to measure the intricate and dynamic aspects of spontaneous psychotic brain activity. These measures can help us identify irregularities and patterns in complex systems. In our study, we utilized fuzzy recurrence plots and sample entropy to evaluate the dynamic characteristics of psychiatric disorders. This assessment focused on understanding the temporal and spatial neural activity patterns, and more specifically, we applied complexity measures to investigate the functional connectivity within the psychotic brain. This involves understanding how different brain regions synchronize their activity, and complexity measures can reveal the patterns of these connections. It provides a means to understand how different brain regions interact and communicate under resting-state abnormal conditions. This study offers evidence demonstrating that fuzzy recurrence plots can serve as descriptors for functional connectivity and discusses their relevance to sample entropy in the context of the psychotic brain. In summary, complexity measures offer valuable insights that enrich our comprehension of atypical brain activity and the complexities present in the psychotic brain^1^.

## 1. INTRODUCTION

Comprehending the computational mechanisms that drive the intricate, nonlinear dynamics within the human brain is an ongoing and challenging scientific pursuit, and the evaluation of complexity in resting-state functional magnetic resonance imaging (fMRI) signals, specifically employing entropy-based techniques, has garnered considerable interest. This complexity is demonstrated by the behavioral traits of the brain, where its components participate in intricate interactions guided by principles of neural computation. These interactions result in non-linear behaviors, collective dynamics, hierarchical arrangements, and emergent phenomena [1–3].

Complexity measures primarily encompass predictability and regularity dimensions, and predictability assesses how well brain activity can be predicted, whereas regularity examines to identify recurrent patterns or order in the data. In general, predictability can manifest as spatial measures (such as the *higuchi fractal dimension* [4], *detrended fluctuation analysis* [5], among others) and temporal measures (including *correlation dimension* [6], *largest lyapunov exponent* [7], and others). Regularity can be further categorized into monoscale metrics (e.g., *approximate entropy* [8], *sample entropy* [9], *lempel-ziv complexity* [10], and more) and multiscale metrics (such as *multiscale entropy* [11], *multivariate multiscale entropy* [12], and others).

In this study, our objective is to quantitatively assess the intricacies of both spontaneous and psychotic brain activity by conducting a thorough analysis of the temporal patterns present in fMRI signals. We explored complexity measures that provided us with the means to assess intricate spatial and temporal patterns in brain activities. These measures have the capability to capture the nonlinear dynamics inherent in spontaneous brain activity and can even be employed for the differentiation of psychotic traits.

## 2. THEORETICAL ANALYSIS

### 2.1. Predictability in Spontaneous Brain Activity

***Recurrence Plots***, constitute a sophisticated method for the visualization of multivariate nonlinear data, with specific application in the context of brain activity [13]. It encapsulates data representation as a binary matrix, with individual elements denoting the recurrences of data states or phases at distinct time points. To illustrate, the concept of recurrence plots can be mathematically defined as follows:

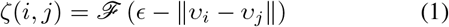

Where *ϵ* serves as a parameter signifying a threshold used to delineate the degree of similarity or dissimilarity between pairs of data points (*υ*_*i*_, *υ*_*j*_). The function *ℱ* is represented as a unit step function, and it yields a value of 0 or 1 depending on whether the condition (*ϵ* − |*υ*_*i*_ − *υ*_*j*_|) *<* 0 holds true or false, respectively.

A recurrence plot is characterized as a binary symmetrical matrix, as denoted by Equation 1. This matrix serves the purpose of visually representing instances in time where the trajectory coincides (depicted as black dots) or diverges (depicted as white dots). To address the challenges posed by establishing a similarity cutoff threshold and the binary nature of recurrence representation in RP, an innovative approach was introduced in the form of a fuzzy recurrence plot (FRP) [14]. In the phase space, the FRP showcases the recurrent behavior of the trajectory through a grayscale image, with values spanning the interval [**0, 1**]. In this representation, black corresponds to **0**, white to **1**, and varying shades of gray signify other values.

The generation of an FRP is achieved by applying the fuzzy c-means algorithm [15], which divides the dataset **X** into a collection of clusters, each denoted as *c* = 3. This algorithm assigns a fuzzy membership grade, symbolized as *ζ*_*ij*_ and taking values within the range of [**0, 1**] (*ζ*_*i,j*_ ∈ [**0, 1**]), to each data point **x**_*i*_, where *i* = 1, 2, …, *k*, concerning its association with each cluster center **v**_*j*_, where *j* = 1, 2, …, *c*. The FRP method incorporates the following properties:

- Reflexivity

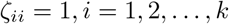
- Symmetry

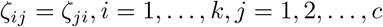
- Transitivity

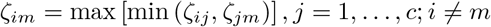

Consequently, a FRP ∈ ℝ^*k×k*^ is formally characterized as a square grayscale image,

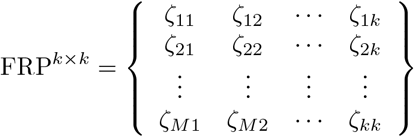

The fMRI signal can be effectively converted into a spatial domain referred to as the dynamical system phase space, and the individual elements within this phase space are termed dynamical system points. The utility of this transformation lies in its capacity to unveil inherent properties within intricate data sets, often surpassing the insights attainable from the original time series itself [13, 14]. Consequently, the selection of the embedding dimension denoted as *e* assumes significance, as it serves to delineate the primary dimensions relevant to the neural signals. Furthermore, determining the time delay parameter, *τ*, and the number of clusters, *c*, within the time series is crucial, as these choices aim to maximize the discernment of nonlinear dynamic features.

### 2.2. Regularity in Spontaneous Brain Activity

***Brain Entropy***, in the context of dynamic systems such as the primate brain, represents the rate at which information is produced. Approximate entropy (ApEn) and sample entropy (SampEn) are two complexity measures closely associated with entropy. They estimate the complexity of a system based on a known prior probabilistic distribution, and they have been extensively utilized in time series analysis [9, 16]. These estimators are valuable for evaluating the complexity of spontaneous brain activity signals. ApEn assesses similarity in time series data, whereas SampEn does not account for self-similar patterns, a feature in ApEn. Furthermore, SampEn offers two distinct advantages over ApEn: it is independent of data length and is comparatively easier to implement.

For an fMRI signal of *𝒩* points, *U* (*i*) (1 ≤ *i* ≤ *𝒩*), informally, given *𝒩* =(*u*_1_, *u*_2_, *u*_3_, ·· ·, *u*_*𝒩*_) points with a constant time interval *t*, the family of statistics *ApEn* (*m, r, 𝒩*) is approximately equal to the negative average natural logarithm of the conditional probability that two sequences that are similar for *m* points remain similar, that is, within a tolerance *r* (usually refers to the distance to consider two data points as similar. and default will set to 0.2 × *std*_*U*(*i*)_, where *std* indicates standard deviation), at the next point. Thus, a low value of ApEn reflects a high degree of regularity. The parameters *m, r*, and *𝒩* must be fixed for each calculation, where *e* for embedding dimension, tolerance *r*, and number of points *𝒩*.

Let’s define a template vector with length *l*, then we have *U*_*l*_(*i*) = {*u*_*i*_, *u*_*i*+1_, *u*_*i*+2_, ···, *u*_*i*+*l*−1_}, and distance function, *𝒟* [*U*_*l*_(*i*), *U*_*l*_(*j*)], where *i* ≠ *j*. Then, we can define the sample entropy as follows,

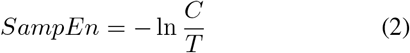

where *C* is the number of template vector pairs having *𝒟*[*U*_*l*+1_(*i*), *U*_*l*+1_(*j*)] *< r*, and *T* is the number of template vector pairs having *𝒟*[*U*_*l*_(*i*), *U*_*l*_(*j*)] *< r*, and considering properties of SampEn [9], the value of SampEn will always be zeros or positive.

## 3. RESTING STATE FMRI DATASET

### 3.1.. Bipolar and Schizophrenia Consortium for Parsing Intermediate Phenotypes (BSNIP) Dataset

A cohort comprising 471 participants was chosen from the Bipolar and Schizophrenia Network for Intermediate Phenotypes (BSNIP) consortium [17]. Within this cohort, 288 individuals were representative of those affected by typical Schizophrenia (SZ), and the remaining 183 had received diagnoses of Bipolar Disorder (BP).

## 4. RESULTS

In order to investigate how complexity metrics might be utilized to analyze brain diseases, a total of 105 independent component networks (ICNs) were identified through the pre-processing of fMRI data, employing spatially constrained independent component analysis (ICA). Concurrently, we developed a brain network template, employing a multi-spatial-scale framework that incorporates 105 ICNs. These ICNs can be categorized into 6 intrinsic brain network domains, as depicted in Fig. 1, and were derived from a substantial dataset encompassing more than 100*K* participants. The average fMRI signal (*ℬ*_*SZ/BP*_) cross subject and ICNs is 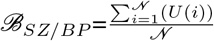, *𝒩* = 105.

**Fig. 1:**
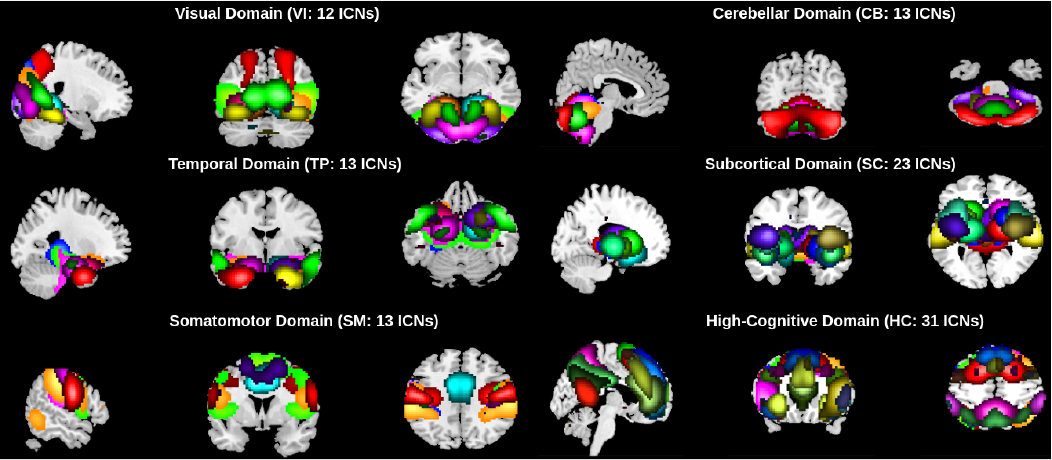
Total of 105 intrinsic brain networks have been systematically classified into six distinct groupings, which are designated as follows: Visual Domain (12 ICNs), Cerebellar Domain (13 ICNs), Temporal Domain (13 ICNs), Subcortical Domain (23 ICNs), Somatomotor Domain (13 ICNs), and High-Cognitive Domain (31 ICNs).

The nonlinear dynamic of the spatial pattern of spontaneous psychotic brain activity was present in Fig. 2 (top), and related optimization of complexity parameters, i.e., *e, τ*, and *r*, based on minimum *Mutual Information* was shown in Fig. 2 (left side), and then low-dimensional manifold attractors were estimated based on optimized parameters, and nonlinear dynamic properties of spontaneous psychotic brain activity can be tracked from the attractor and followed by the FRP spatial pattern (Fig. 2 middle and right sides), and it gives us a window to estimate resting-state functional connectivity while considering the nonlinear dynamic of psychotic brain activity.

**Fig. 2:**
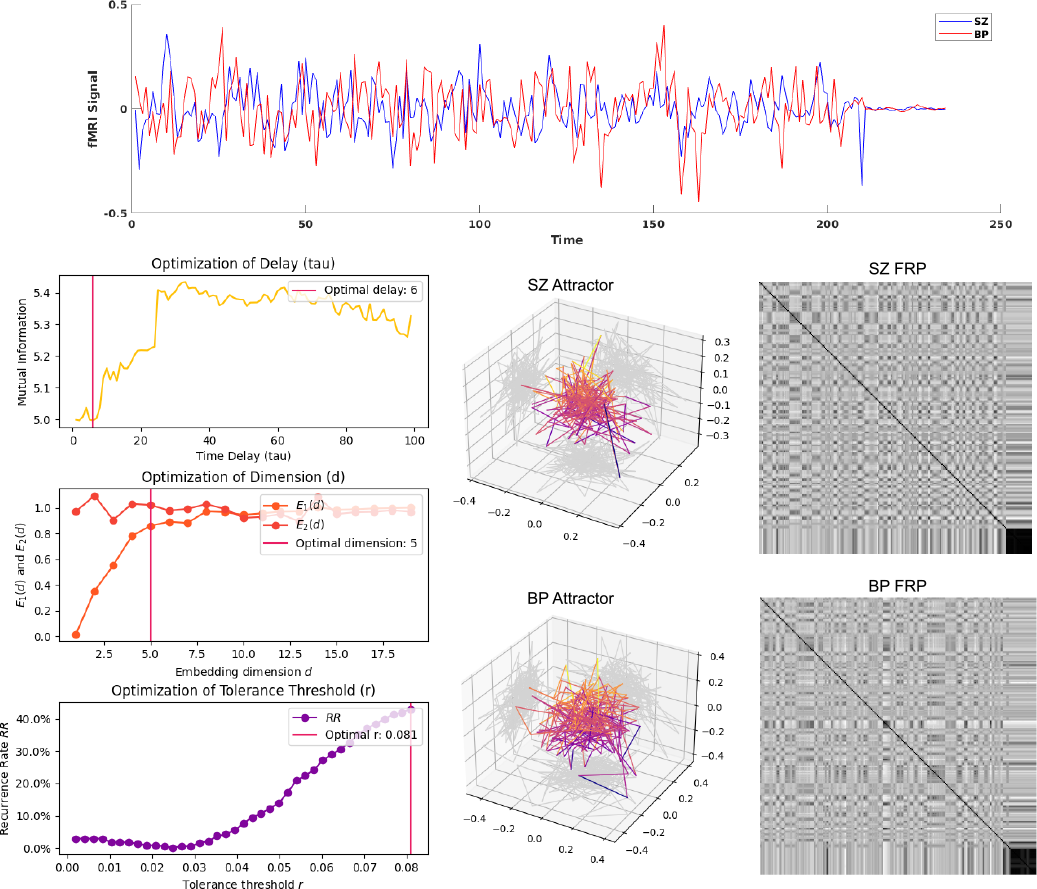
Nonlinear dynamic estimation with attractors and FRP for the BSNIP dataset. The psychotic brain activity is shown at the top. The optimized complexity parameter estimations are shown in the bottom left side (optimized parameter: delay *τ* = 6, embedding dimension *e* = 5, tolerance *r* = 0.081), the related attractors for the psychotic brain are present in the bottom middle, and FRP (optimized parameter: *e* = 3, *τ* = 1, *c* = 3) are presented in the bottom right side.

Then we applied sample entropy to quantify the regularity and unpredictability of fluctuations in psychotic brain signals. As shown in Fig. 3, following the way to optimize complexity parameters, we selected the parameters *e* = 1 and *r* = 0.32 for SZ and BP studies. In Fig. 3, a low sample entropy implies that psychotic brain activity is deterministic, and a large value shows that it is random. Therefore, we showed that BP tends to be more random in brain activity than SZ, which may help us to distinguish between SZ and BP. Moreover, the main topology joint similarity and differences functional brain regions based on *Euclidean Distance, d* = |*SampEn*_*SZ*_ − *SampEn*_*BP*_|, was estimated from sample entropy, *SampEn*_*SZ*_ and *SampEn*_*BP*_, then the joint similarities were 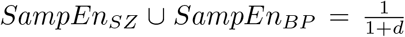, and the difference will be *SampEn*_*SZ*_ ∩ *SampEn*_*BP*_ =1 − (*SampEn*_*SZ*_ ∪ *SampEn*_*BP*_), and the largest differences are mainly present in the visual domain, temporal domain, somatomotor domain, and high-cognitive domain, and we hypothesized that these functional brain regions could be relevant in identifying SZ and BP patients.

**Fig. 3:**
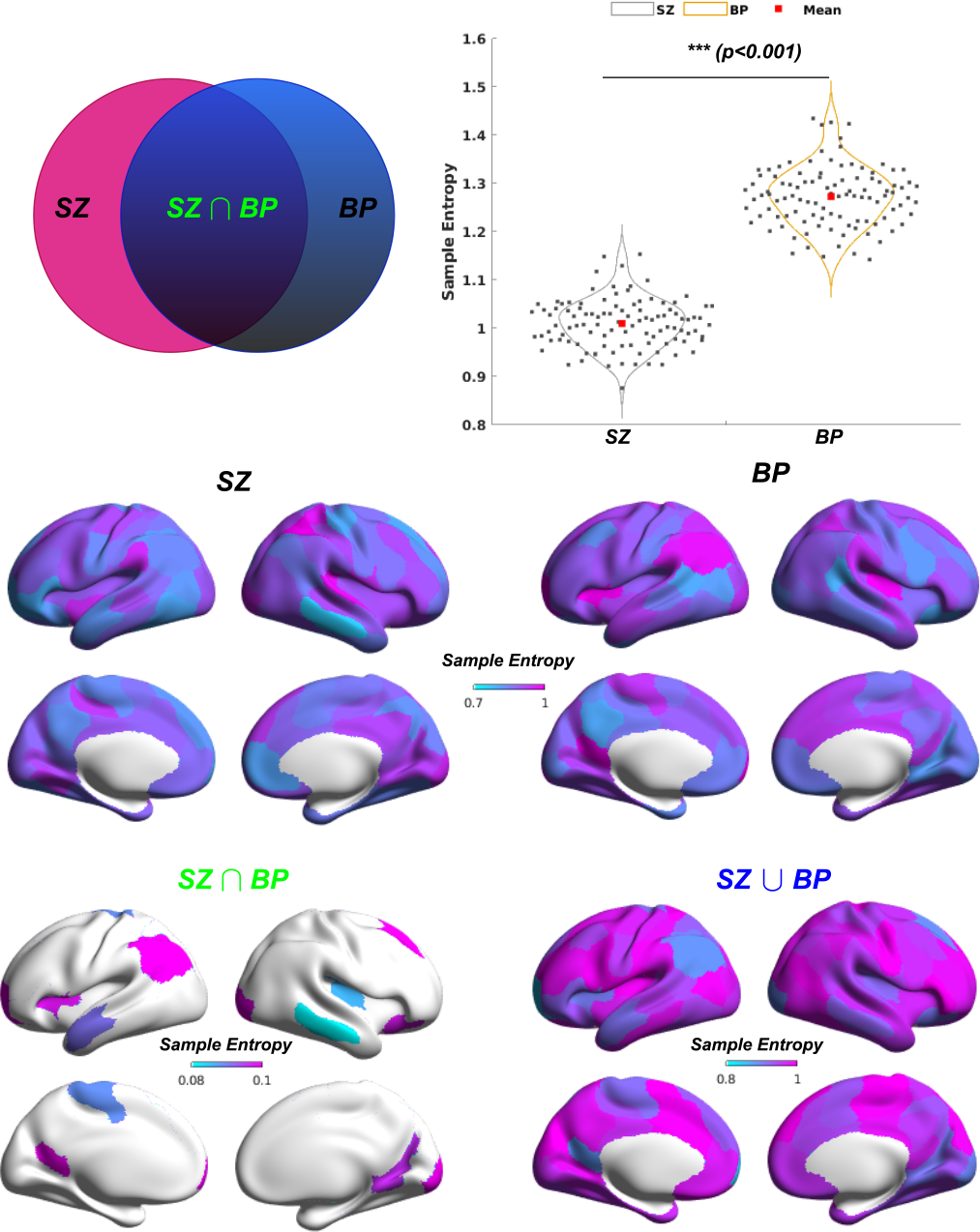
Sample entropy analysis was conducted on the psychotic brain. The resulting sample entropy values for SZ and BP patients were projected onto the standard brain surface (*fs*_*LR*.32*k*), revealing notable topological joint and distinctions considering SZ and BP traits are hard to separate. Related statistical analyses demonstrated significant sample entropy differences expressed between SZ and BP.

## 5. DISCUSSION

Complexity measures show great potential for quantifying brain activity and may offer assistance in diagnosing psychotic disorders. This marks the initial exploration of the topic, with the next phase involving a more organized investigation into the synchronization of neural activity across various cognitive states. Simultaneously, there can be focused efforts on pinpointing potential biomarkers for the accurate diagnosis of brain disorders using complexity measures.

## 6. ACKNOWLEDGEMENTS

This work was supported by NSF grant 2112455, and NIH grants R01MH123610 and R01MH119251.

## Footnotes

1 IEEE Southwest Symposium on Image Analysis and Interpretation (SSIAI 2024)

